# mtx-COBRA: Subcellular localization prediction for bacterial proteins

**DOI:** 10.1101/2023.10.04.560913

**Authors:** Isha Arora, Arkadij Kummer, Hao Zhou, Mihaela Gadjeva, Eric Ma, Gwo-Yu Chuang, Edison Ong

## Abstract

**Background:** Bacteria can have beneficial effects on our health and environment; however, many are responsible for serious infectious diseases, warranting the need for vaccines against such pathogens. Bioinformatic and experimental technologies are crucial for the development of vaccines. The vaccine design pipeline requires identification of bacteria-specific antigens that can be recognized and induce a response by the immune system upon infection. Immune system recognition is influenced by the location of a protein. Methods have been developed to determine the subcellular localization (SCL) of proteins in prokaryotes and eukaryotes. Bioinformatic tools such as PSORTb can be employed to determine SCL of proteins, which would be tedious to perform experimentally. Unfortunately, PSORTb often predicts many proteins as having an “Unknown” SCL, reducing the number of antigens to evaluate as potential vaccine targets.

**Method:** We present a new pipeline called sub<u>C</u>ellular l<u>O</u>calization prediction for <u>B</u>acte<u>R</u>i<u>A</u>l Proteins (mtx-COBRA). mtx-COBRA uses Meta’s protein language model, Evolutionary Scale Modeling, combined with an Extreme Gradient Boosting machine learning model to identify SCL of bacterial proteins based on amino acid sequence. This pipeline is trained on a curated dataset that combines data from UniProt and the publicly available ePSORTdb dataset.

**Results:** Using benchmarking analyses, nested 5-fold cross-validation, and leave-one-pathogen-out methods, followed by testing on the held-out dataset, we show that our pipeline predicts the SCL of bacterial proteins more accurately than PSORTb.

**Conclusions:** mtx-COBRA provides an accessible pipeline with greater efficiency to classify bacterial proteins with currently “Unknown” SCLs than existing bioinformatic and experimental methods.

## 1. Introduction

Bacteria are widely present in the environment and are necessary to maintain homeostasis. They play important roles in the upkeep of our bodily functions, ranging from digestive to reproductive processes [1]. However, some bacteria are pathogenic, causing serious diseases such as bacterial pneumonia (*Streptococcus pneumoniae*) [2] and meningitis (meningococcal bacteria) [3]. Within 2019 alone, there were 7.7 million deaths associated with 33 common bacterial infections [4]. Recently, there has been an increase in bacterial antibiotic resistance for multiple reasons including overuse, inappropriate prescriptions, and widespread agricultural use of antibiotics [5]. This era of increasing antibiotic resistance is a global concern and has even been declared as a potentially debilitating crisis by the World Health Organization (WHO) [6]. A group of bacteria known as the carbapenem-resistant Enterobacteriaceae are resistant to nearly all contemporary antibiotics, resulting in a survival rate of only 50% for patients hospitalized following infection [7]. Furthermore, bacteria that have gained antibiotic resistance are associated with 23,000 deaths per year within the United States, with similar statistics in Europe [8].

An alternative strategy to antibiotics is vaccination, which aids in the prevention of infectious diseases and reduces the need for antibiotics and other therapies [9]. Currently, there are multiple Food and Drug Administration–approved vaccinations for various bacterial infections, including Typhim Vi and Vivotif for typhoid, MenQuafdi and Trumenba for meningococcal disease, and Boostrix and Adacel for tetanus, diphtheria, and pertussis [10–15]. Some vaccines, such as the bacilli Calmette-Guérin vaccine for tuberculosis, confer suboptimal protection [16]. The need for more effective vaccines is emphasized by the acquisition of multidrug resistant tuberculosis or extensively-drug resistant tuberculosis, both of which present significant challenges in disease treatment. According to the WHO, approximately 470,000 people are infected by multidrug resistant tuberculosis, of which about 180,000 succumb to the infection [17]. Further, there are deadly bacteria against which vaccinations have yet to be developed, such as *Pseudomonas aeruginosa*, *Staphylococcus aureus*, and *Neisseria gonorrhoeae* [18]. In all cases, vaccines are necessary to improve protection against these bacterial infections.

When developing vaccines against bacterial pathogens, it is essential to test the immunogenicity of various antigens *in vivo*. Bacterial proteomes generally have thousands of proteins which cannot be individually tested using wet lab methods because of cost, time, and labor [19]. Hence, bioinformatic reverse vaccinology tools have been increasingly applied to address the need for down selection of antigens [20]. Such a pipeline generally includes comparative genomics, which involves the determination of conserved antigens among various strains of the bacteria and ensures that the antigens are unique to the pathogenic strains, among other steps. An essential part of this pipeline is determining the subcellular localization (SCL) of the various antigens identified, which can help restrict the list of target antigens to those present on the surface or secreted by bacteria. [21] Identifying protein SCL is important because the immune system can more readily recognize and develop an immune response against proteins that are visible to the body, that is, secreted or membrane-bound proteins [22].

Five potential SCLs exist for gram-negative proteins comprising the cytoplasm, cytoplasmic membrane, periplasm, outer membrane, and the extracellular region. Four potential SCLs exist for gram-positive bacteria comprising the cytoplasm, cytoplasmic membrane, cell wall, and the extracellular region [23]. Proteins that are exposed and easily recognized by the human immune system are expressed in the outer membrane or extracellular region of gram-negative bacteria and the cell wall or extracellular region of gram-positive bacteria [24].

SCL can be determined *in vitro* by extracting protein samples from various fractions of the bacteria of interest and digesting these proteins into peptides. The peptides would then be differentially labelled to undergo liquid chromatography-tandem mass spectrometry; the abundance ratios for each SCL can then be calculated. [25] Fluorophore tagging of the protein followed by microscopy is another method used in the determination of SCL *in vitro* [26]. Analytical and labeling techniques are lengthy and expensive, thus, performing such experiments for the thousands of proteins in one bacterium is not reasonably feasible [19]. Therefore, there is a need for bioinformatic methods that can accurately predict the SCL of a protein given its sequence, while simultaneously reducing the cost and time usually required to determine the SCL of a protein or potential antigen.

Multiple bioinformatic methods have been developed for predicting SCLs, including PSORTb, CELLO2GO, and SubLoc, wherein PSORTb is designed specifically for gram-negative and gram-positive bacteria and archaea, and CELLO2GO and SubLoc are designed to predict the SCL for proteins from prokaryotes and eukaryotes [25, 27, 28]. PSORTb uses various support vector machines (SVMs) in combination with a Bayesian network to predict SCL [28], CELLO2GO uses BLAST to obtain gene ontology–annotated homologous sequences which aids its prediction of SCL [25], and SubLoc uses an SVM for SCL predictions [27]. In a benchmarking study of these tools, PSORTb was reported to have higher precision and recall scores than CELLO2GO for both gram-negative and gram-positive bacteria [28].

PSORTb has been the widely accepted program for SCL predictions. The authors of PSORTb also released two accompanying datasets called ePSORTdb and cPSORTdb; the former contains proteins and their experimentally determined SCLs, and the latter contains proteins along with their predicted SCLs determined by PSORTb [28]. Despite the benefits of PSORTb, when used to predict protein sequences that are not included in ePSORTdb, PSORTb returns many proteins labeled with an “Unknown” SCL prediction. This prevents many antigens from being evaluated as potential vaccine candidates when used in the reverse vaccinology pipeline; that is, they may truly be present on the surface of the bacteria, but PSORTb predicts them as “Unknown.”

To overcome this, we propose a new pipeline, sub<u>C</u>ellular l<u>O</u>calization prediction for <u>B</u>acte<u>R</u>i<u>A</u>l Proteins (mtx-COBRA), for SCL predictions that uses Meta’s protein language model, Evolutionary Scale Modeling (ESM) [29] followed by a pre-trained Extreme Gradient Boosting (XGBoost) machine learning (ML) model. Two models were used—one for gram-negative bacteria and the other for gram-positive bacteria. These models were trained on a combination of representative sequences from ePSORTdb and a curated dataset using UniProt data for various gram-negative and gram-positive bacteria, respectively. Nested 5-fold cross-validation (5FCV) and leave-one-pathogen-out (LOPO) were conducted for initial testing of the performance of mtx-COBRA. The performance of mtx-COBRA and PSORTb were then compared and tested on a held-out dataset to select the XGBoost ML model from all considered models.

## 2. Methods

### 2.1. Dataset collection

The project workflow is described in **Fig. 1**. Briefly, three datasets were used to complete the analysis: ePSORTdb, the expanded dataset, and a held-out dataset. The ePSORTdb dataset was one of the datasets used to train PSORTb and is composed of sequences that have experimentally determined SCLs [28, 30]. Version 4 of ePSORTdb was downloaded from https://db.psort.org/downloads and used in training and testing of the models as described below. Protein IDs were passed through the UniProt, UniParc, National Center for Biotechnology Information (NCBI), and Entrez databases using their respective REpresentational State Transfer (REST) application programming interfaces (APIs) and the required FASTA sequences were downloaded [31–34].

**Fig. 1.**
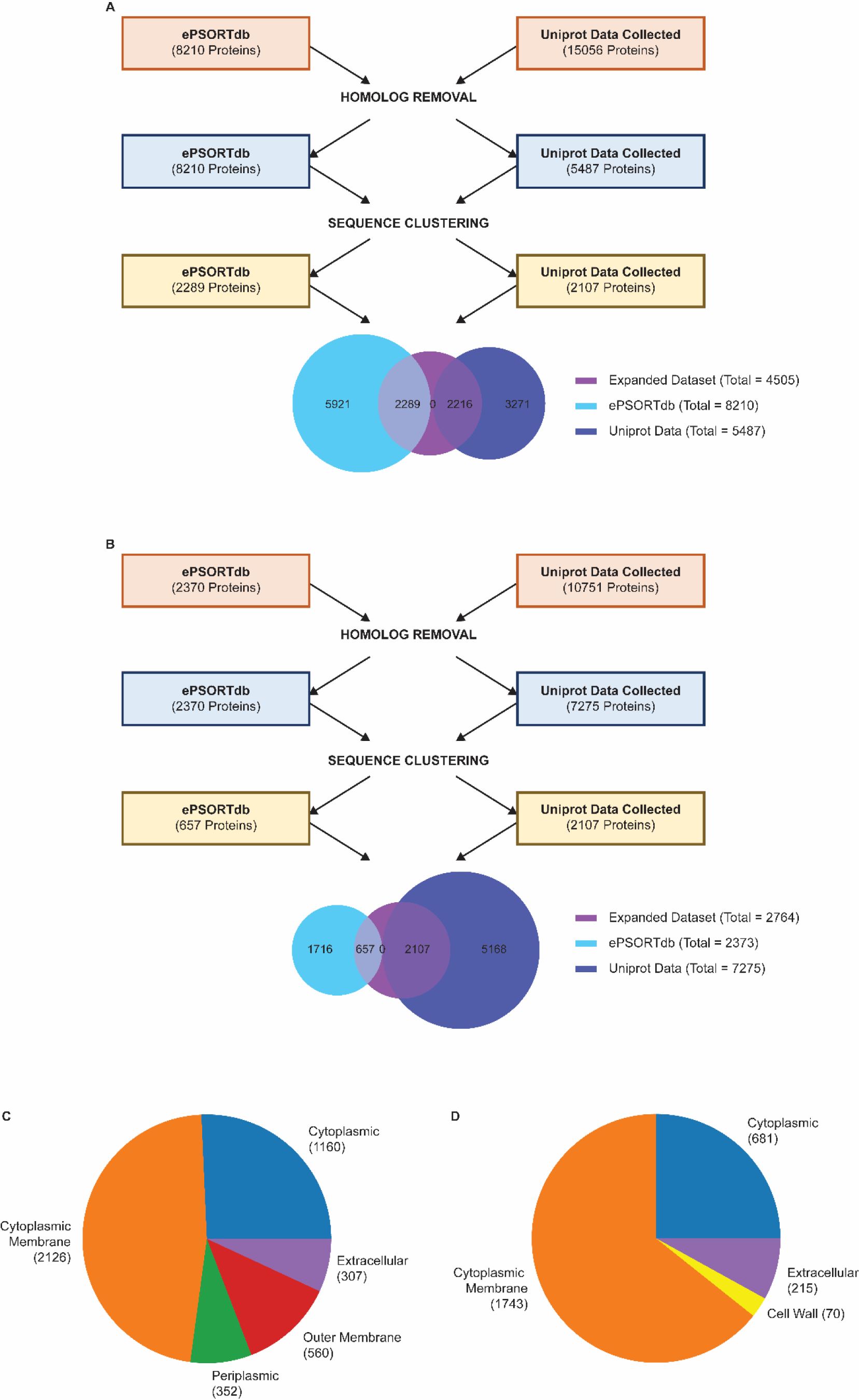
Project workflow. Pipeline depicting the data collection process of proteins in the expanded, ePSORTdb, and UniProt datasets for (A) gram-negative bacteria and (B) gram-positive bacteria. Venn diagram depicting SCLs and the corresponding number of proteins in the expanded (C) gram-negative and (D) gram-positive datasets.

The gram-negative expanded dataset was generated from 8210 proteins from ePSORTdb and 15,056 from UniProt which were downloaded, processed, down-selected, and combined to form a database of 2289 proteins from ePSORTdb and 2216 from UniProt. The proteomes and respective metadata for 13 gram-negative bacteria were selected and downloaded from UniProt using the Proteins REST API [35] and the UniProt REST API. UniProt proteins with SCL evidence beyond “sequence similarity” (ECO:0000250) and “sequence motif match” (ECO:0000259) were included for further evaluation. This process was repeated to create the gram-positive expanded dataset, with 17 gram-positive bacteria from UniProt combined with ePSORTdb to create a final of 657 proteins from ePSORTdb and 2107 proteins from UniProt.

To avoid potential bias in the expanded dataset, duplicated proteins and homologs between ePSORTdb and UniProt were removed using Diamond BLAST [36] with a 30% sequence identity cutoff. Representative sequences from the resultant UniProt data were obtained by using Many-against-Many sequence searching (MMseqs2) with default settings. These sequences were combined with ePSORTdb to obtain the final expanded dataset of 4505 gram-negative bacterial proteins and 2764 gram-positive bacterial proteins (**Fig. 1**).

A similar pipeline was conducted to generate the third, held-out dataset. Initially, 13,368 proteins from 24 new gram-negative bacteria and 22,316 proteins from 94 gram-positive bacteria in the UniProt database were collected. Proteins with evidence numbers ECO:0000250 or ECO:0000259 were not included in the database. Following the same strategy used to generate the expanded dataset, duplicated proteins and homologs were removed, and representative sequences were selected and retained for the final held-out dataset, leaving 1380 proteins in the final gram-negative and 1660 proteins in the final gram-positive held-out datasets.

### 2.2. Overall computational pipeline

The proposed pipeline used the Meta ESM protein language model [29], followed by an ML model to predict the SCLs of the various proteins from their amino acid sequences. To train the ML model, all protein sequences in the expanded and held-out datasets were passed through ESM to obtain a matrix of features. Then these features and the SCLs encoded as integer labels were fed into different ML models for training and evaluation. The optimal ML model for this pipeline was determined using XGBoost, SVM, k-nearest neighbors (KNN), neural network, naïve Bayes, random forest, and logistic regression models which were tested using the nested 5FCV and LOPO benchmarking methods (**Fig. 2**).

**Fig. 2.**
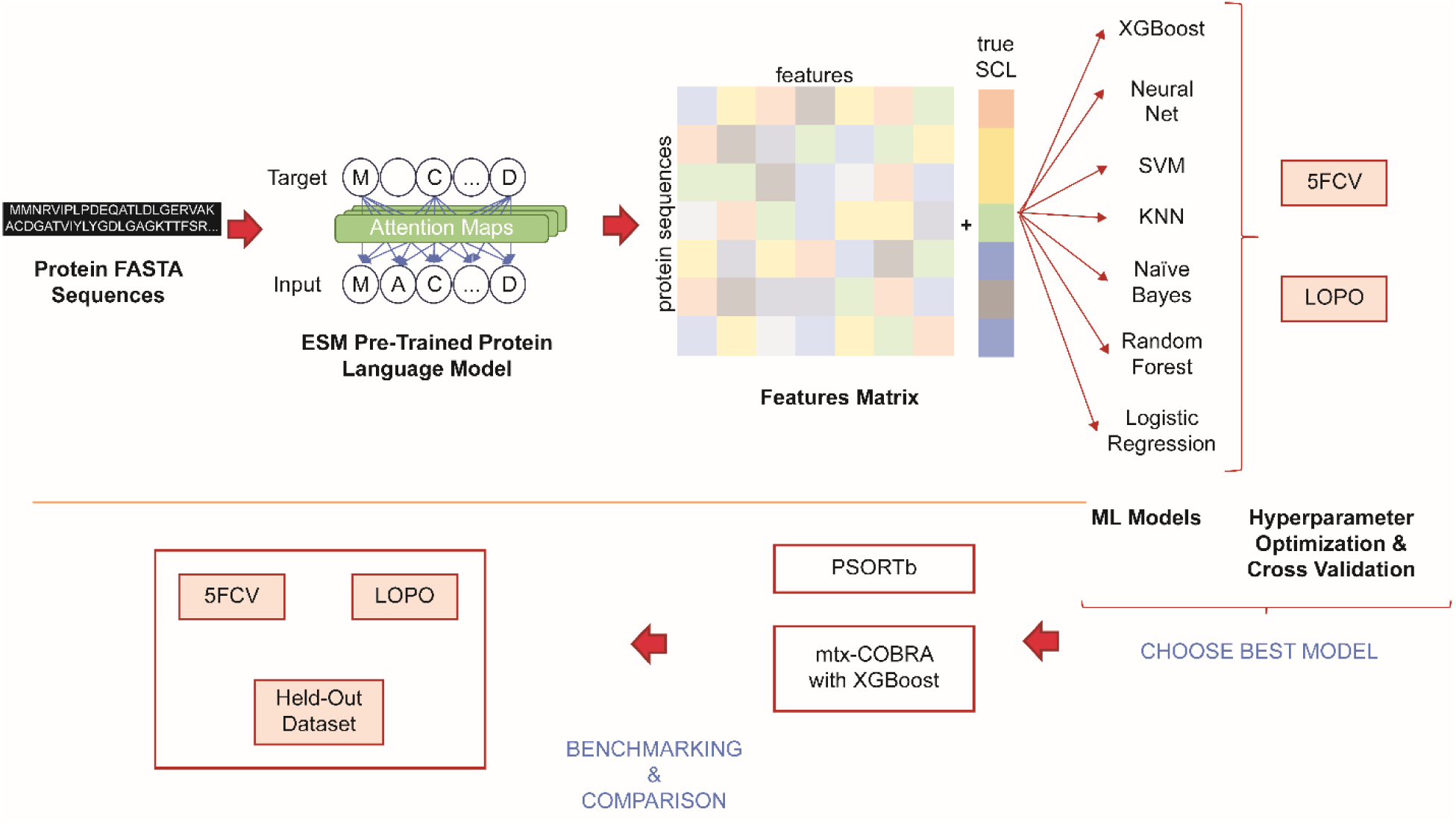
Visual depiction of our pipeline to predict SCL.

To obtain a features matrix, protein FASTA sequences were passed through the pre-trained ESM protein language model by Meta [29]. This features matrix and the corresponding integer labels for the true SCLs were passed to the ML model (such as XGBoost) to train and test on the ML model.

### 2.3. Nested 5FCV

Initially, the expanded dataset was used for training and testing of the first round of benchmarking. The expanded dataset was passed through ESM to obtain the required features matrix that was then split into the training and testing datasets. For each ML model tested in the pipeline, the training dataset underwent two loops involving stratified shuffle splits. The outer loop divided the data into five folds and was used to determine the best model for our pipeline. For each of these five folds, an inner loop was run wherein another stratified shuffle split where K = 5 was conducted to ensure hyperparameter optimization for the model. Once hyperparameter optimization was conducted, the selected hyperparameters and the training data from the stratified split in the outer loop were used to train the ML model. The model’s performance was tested on the data from the stratified shuffle split in the outer loop. The performance of the pipeline for a particular dataset and ML model was assessed using the averaged classification metrics including precision, recall, F1 score, and Matthew’s correlation coefficient (MCC) scores obtained from each fold in the outer loop. In these analyses, the precision, recall, and F1 scores were adjusted for MCC score. The model was retrained on the SCL data for the binary classifications and then the predictions were compared with the true values to recalculate the precision, recall, F1, and MCC scores.

A precision-recall curve was plotted for each ML model, fold, and SCL when the model was trained and tested on the expanded dataset. The results from all the SCLs were concatenated and used to plot an average precision recall curve for each fold and model. This process was repeated with ePSORTdb.

The true SCLs for the ML models of the expanded dataset and ePSORTdb were converted into a binary format wherein the locations were identified as surface exposed or non-surface exposed. That is, for gram-negative bacteria, the outer membrane and extracellular locations were categorized as surface exposed, whereas the cytoplasmic, cytoplasmic membrane, and periplasmic locations were categorized as non–surface exposed. As for gram-positive bacteria, the cell wall and extracellular locations were categorized as surface exposed, and the cytoplasmic and cytoplasmic membrane locations were categorized as non–surface exposed. After converting the SCLs to this binary format, 5FCV was conducted as described. The ML model was retrained on ESM features matrix for each fold of the training dataset and the corresponding truth values for the binary SCLs. The binary SCLs were predicted for each sequence in the test dataset, and the precision, recall, F1, and MCC scores were recalculated.

### 2.4. LOPO

The LOPO cross-validation method trained the model on the data corresponding to all but one bacterium (training data) and testing the performance of the model on that single organism. To allow for fair testing methodologies, the expanded dataset was filtered to identify which bacteria have proteins expressed in all respective SCLs. This left 13 gram-negative bacteria with 2989 proteins and 10 gram-positive bacteria with 1207 proteins. The taxonomy trees as taken from NCBI’s Common Taxonomy Tree [37, 38] are provided in **Supplementary Fig. 1.**

Each of the bacteria in the LOPO dataset was kept for testing, and thus, the rest of the training dataset underwent a stratified shuffle split with K = 5 to assist in hyperparameter optimization. These selected parameters were first used to train the model on the updated training dataset without the selected bacterium and then were tested on the left-out bacterium. The precision, recall, F1, and MCC scores were calculated for all gram-negative and gram-positive bacteria included in either dataset; the values from all rounds of testing were averaged to determine the performance of the model when trained and tested on the expanded dataset using the LOPO method. A precision-recall curve was plotted for each MCC, ML model, bacterium, and SCL. The results from all the SCLs were concatenated, and an average precision recall curve was plotted for each fold and model.

Similar to 5FCV, the true SCLs for all proteins in the dataset were converted into a binary format. The model underwent LOPO cross-validation over the organisms as described above. For each of the folds, the ESM features for each sequence in the training set and the corresponding binary SCLs were used to retrain the ML model, and the binary SCLs were predicted for the proteins corresponding to the organism in the test set. The precision, recall, F1, and MCC scores were recalculated for each fold and averaged over all organisms.

### 2.5. Benchmarking of mtx-COBRA

The ML model with the best performance, termed mtx-COBRA, was trained on the entire expanded dataset using XGBoost and benchmarked against PSORTb. For this purpose, we applied 5FCV and LOPO with the additional held-out dataset to assess the performance between the two models. For all three evaluations (5FCV, LOPO, and held-out), the predicted SCLs from mtx-COBRA and PSORTb were compared with the true SCLs, and the precision, recall, F1, and MCC scores were calculated. To determine the best ML model, the classification metrics were compared with those obtained from the analyses of different ML models within the pipeline.

In the calculations of precision, recall, F1, and MCC scores, each SCL was set as its own class. Specifically, for PSORTb, the “Unknown” label was considered a separate class, and when a protein was labeled as having an “Unknown” SCL, the SCL was considered to be incorrect.

## 3. Results

### 3.1. Nested 5FCV benchmarking

The performance of each ML model for the first round of benchmarking was assessed using 5FCV. From this analysis, the sample-weighted precision, recall, F1, and MCC scores were recorded for the multi-class and binary classifications on the expanded datasets for the gram-negative and gram-positive bacteria in **Tables 1 and 2**, respectively. The top three performing models in our pipeline were XGBoost, neural network, and SVM; KNN, logistic regression, and random forest performed poorly in comparison. When run on the expanded dataset, XGBoost performed the best out of all the ML models, demonstrating the greatest precision, recall, F1, and MCC scores for both the multi-class and binary classifications. To further assess the performance of each ML model, we applied the same 5FCV on models trained with the ePSORTdb dataset. The performance was comparable to the gram-negative and gram-positive bacteria expanded datasets (**Supplementary Tables 1 and 2**, respectively).

**Table 1.**
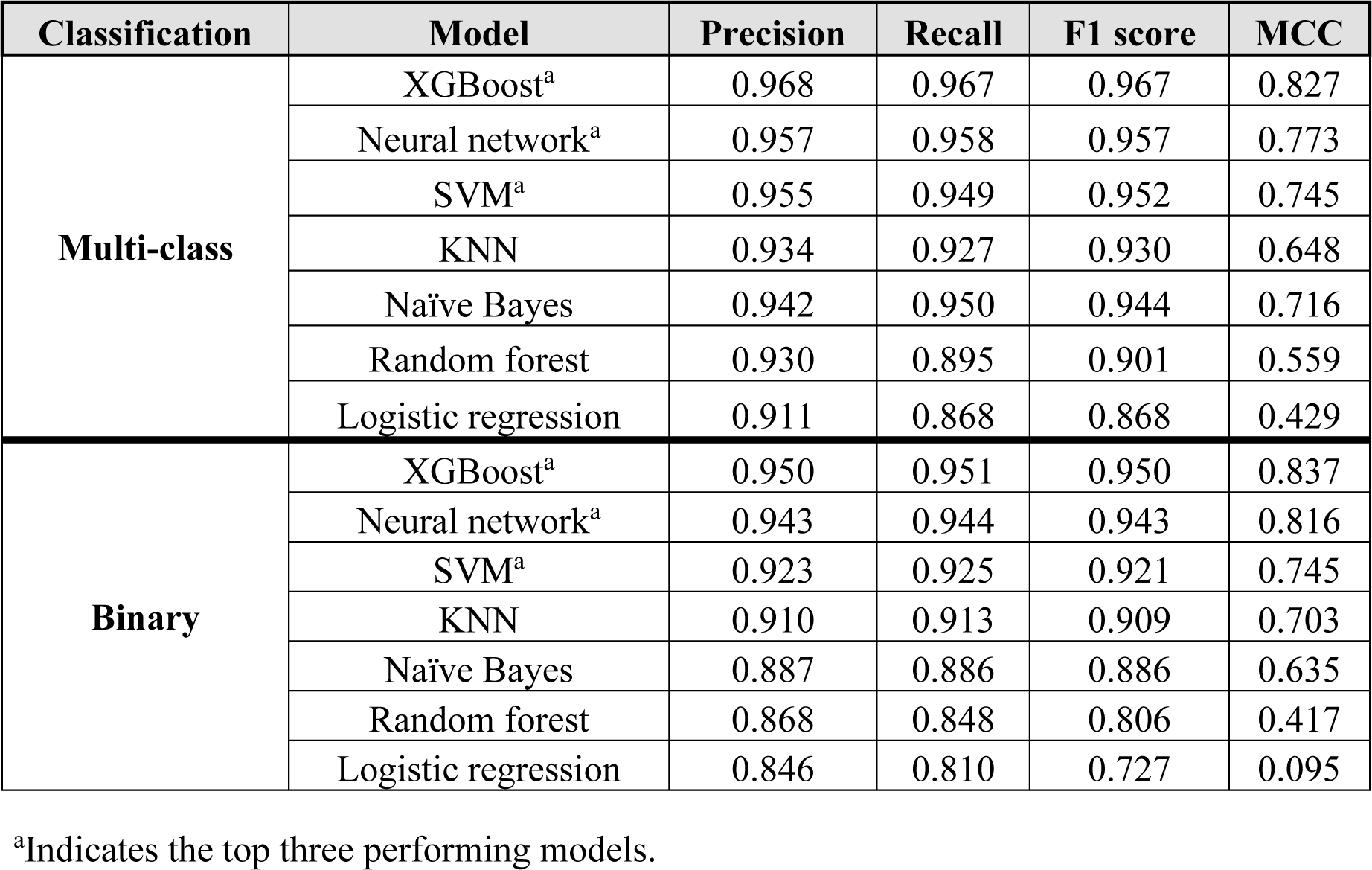
Average of precision, recall, F1, and MCC scores for each model run on the gram-negative bacteria expanded dataset using the 5FCV method.

| Classification | Model | Precision | Recall | F1 score | MCC |
| --- | --- | --- | --- | --- | --- |
| Multi-class | XGBoost<sup>a</sup> | 0.968 | 0.967 | 0.967 | 0.827 |
|  | Neural network<sup>a</sup> | 0.957 | 0.958 | 0.957 | 0.773 |
|  | SVM<sup>a</sup> | 0.955 | 0.949 | 0.952 | 0.745 |
|  | KNN | 0.934 | 0.927 | 0.930 | 0.648 |
|  | Naïve Bayes | 0.942 | 0.950 | 0.944 | 0.716 |
|  | Random forest | 0.930 | 0.895 | 0.901 | 0.559 |
|  | Logistic regression | 0.911 | 0.868 | 0.868 | 0.429 |
| Binary | XGBoost<sup>a</sup> | 0.950 | 0.951 | 0.950 | 0.837 |
|  | Neural network<sup>a</sup> | 0.943 | 0.944 | 0.943 | 0.816 |
|  | SVM<sup>a</sup> | 0.923 | 0.925 | 0.921 | 0.745 |
|  | KNN | 0.910 | 0.913 | 0.909 | 0.703 |
|  | Naïve Bayes | 0.887 | 0.886 | 0.886 | 0.635 |
|  | Random forest | 0.868 | 0.848 | 0.806 | 0.417 |
|  | Logistic regression | 0.846 | 0.810 | 0.727 | 0.095 |

**Table 2.** Average of precision, recall, F1, and MCC scores for each model run on the gram-positive bacteria expanded dataset using 5FCV method.

| Classification | Model | Precision | Recall | F1 score | MCC |
| --- | --- | --- | --- | --- | --- |
| Multi-class | XGBoost<sup>a</sup> | 0.981 | 0.981 | 0.981 | 0.895 |
|  | Neural network<sup>a</sup> | 0.978 | 0.980 | 0.979 | 0.882 |
|  | SVM<sup>a</sup> | 0.977 | 0.977 | 0.977 | 0.873 |
|  | KNN | 0.943 | 0.922 | 0.930 | 0.651 |
|  | Naïve Bayes | 0.949 | 0.964 | 0.954 | 0.764 |
|  | Random forest | 0.952 | 0.899 | 0.912 | 0.612 |
|  | Logistic regression | 0.956 | 0.920 | 0.932 | 0.680 |
| Binary | XGBoost<sup>a</sup> | 0.969 | 0.970 | 0.969 | 0.832 |
|  | Neural network<sup>a</sup> | 0.976 | 0.976 | 0.976 | 0.871 |
|  | SVM<sup>a</sup> | 0.968 | 0.969 | 0.968 | 0.829 |
|  | KNN | 0.944 | 0.947 | 0.943 | 0.692 |
|  | Naïve Bayes | 0.946 | 0.941 | 0.941 | 0.702 |
|  | Random forest | 0.865 | 0.897 | 0.850 | 0.096 |
|  | Logistic regression | 0.909 | 0.903 | 0.864 | 0.262 |

<sup>a</sup>Indicates the top three performing models.

The average precision-recall curve over all folds for each model was plotted to further quantify the performance of the ML models on the expanded dataset (**Fig. 3**). Additionally, for each model, the precision-recall curves for each fold were plotted individually for the gram-negative and gram-positive bacteria (**Supplementary Figs. S2 and S3**, respectively).

**Fig. 3.**
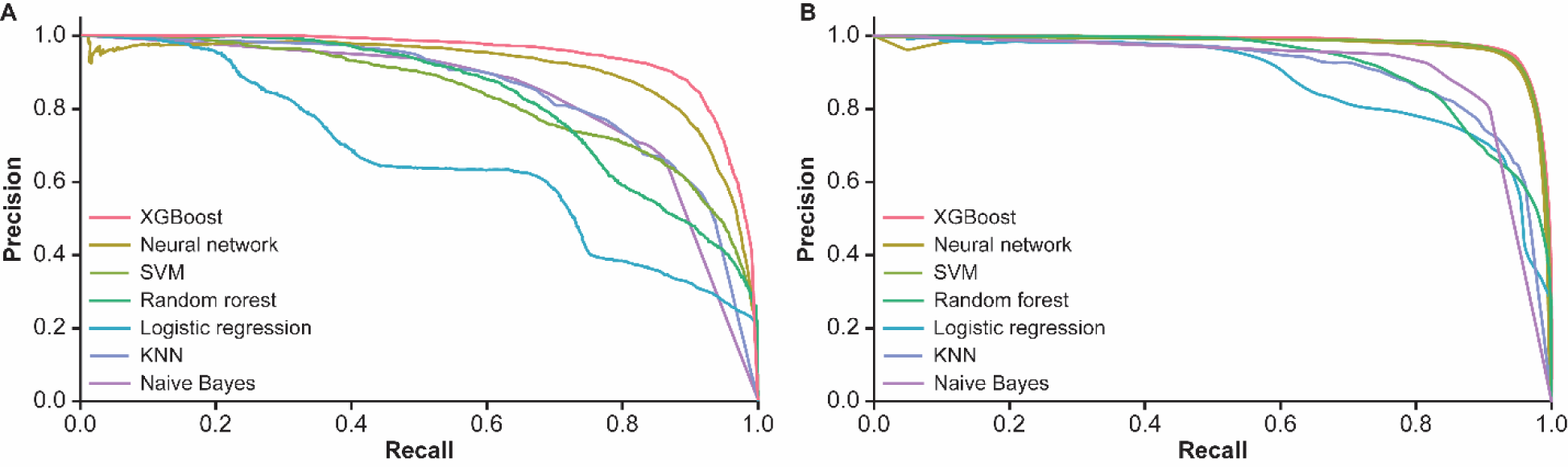
Average precision-recall curves over the five folds for each of the ML models in nested 5FCV (multi-class classification) for (A) the 13 gram-negative bacteria and (B) 10 gram-positive bacteria.

Our results demonstrated that XGBoost performed well in both the multi-class and binary classification cases and was the best-performing model using both classification strategies for gram-negative bacteria and the multi-class classification for gram-positive bacteria. The neural network, with two layers of 150 nodes each, was the best-performing model in the binary case for gram-positive bacteria.

The precision-recall curves and the area under the curve (AUC) for each of the folds for XGBoost and neural network further clarified successful performance of both models.

The AUC was 0.97 and 0.96 for XGBoost and neural network, respectively; thus, the XGBoost model performed slightly better in the multi-class case.

After being trained on the respective gram-negative and gram-positive bacteria expanded datasets, the models were also trained and tested on ePSORTdb and the scores were recalculated (**Supplementary Tables S3 and S4**). The performance of the pipeline was similar when trained on either the ePSORTdb or the expanded dataset.

### 3.2. LOPO benchmarking

LOPO cross-validation method was used for the second round of multi-class and binary classification benchmarking on the expanded datasets for both gram-negative and gram-positive bacteria. XGBoost, neural network, and SVM were the top performing models for both gram-negative and gram-positive bacteria (**Tables 3 and 4**, respectively). XGBoost performed the best on the gram-negative expanded dataset and performed well in both the multi-class and binary classification cases of the gram-positive expanded dataset. However, the neural network, with two layers of 150 nodes each, was the best performing model in the multi-class case of the gram-positive expanded dataset. Both models had a similar performance with comparable MCC scores across multi-class and binary classification among the gram-negative and gram-positive expanded datasets.

**Table 3.** Average of precision, recall, F1, and MCC scores for each model run on the gram-negative expanded dataset using LOPO cross-validation method.

| Classification | Model | Precision | Recall | F1 score | MCC |
| --- | --- | --- | --- | --- | --- |
| Multi-class | XGBoost<sup>a</sup> | 0.966 | 0.965 | 0.965 | 0.818 |
|  | Neural network<sup>a</sup> | 0.963 | 0.957 | 0.958 | 0.790 |
|  | SVM<sup>a</sup> | 0.955 | 0.945 | 0.948 | 0.743 |
|  | KNN | 0.880 | 0.862 | 0.856 | 0.351 |
|  | Naïve Bayes | 0.944 | 0.955 | 0.947 | 0.731 |
|  | Random forest | 0.931 | 0.897 | 0.898 | 0.572 |
|  | Logistic regression | 0.916 | 0.872 | 0.869 | 0.450 |
| Binary | XGBoost<sup>a</sup> | 0.945 | 0.941 | 0.939 | 0.797 |
|  | Neural network<sup>a</sup> | 0.937 | 0.932 | 0.932 | 0.777 |
|  | SVM<sup>a</sup> | 0.947 | 0.946 | 0.943 | 0.803 |
|  | KNN | 0.867 | 0.849 | 0.833 | 0.435 |
|  | Naïve Bayes | 0.917 | 0.904 | 0.908 | 0.674 |
|  | Random forest | 0.884 | 0.845 | 0.807 | 0.442 |
|  | Logistic regression | 0.736 | 0.804 | 0.732 | 0.126 |

<sup>a</sup>Indicates the top three performing models.

**Table 4.** Average of precision, recall, F1, and MCC scores for each model run on the gram-positive expanded dataset using LOPO cross-validation method.

| Classification | Model | Precision | Recall | F1 score | MCC |
| --- | --- | --- | --- | --- | --- |
| Multi-class | XGBoost<sup>a</sup> | 0.968 | 0.966 | 0.967 | 0.857 |
|  | Neural network<sup>a</sup> | 0.971 | 0.969 | 0.969 | 0.868 |
|  | SVM<sup>a</sup> | 0.969 | 0.968 | 0.968 | 0.860 |
|  | KNN | 0.879 | 0.827 | 0.827 | 0.376 |
|  | Naïve Bayes | 0.927 | 0.940 | 0.930 | 0.711 |
|  | Random forest | 0.951 | 0.931 | 0.939 | 0.750 |
|  | Logistic regression | 0.935 | 0.890 | 0.902 | 0.635 |
| Binary | XGBoost<sup>a</sup> | 0.967 | 0.967 | 0.965 | 0.800 |
|  | Neural network<sup>a</sup> | 0.955 | 0.953 | 0.951 | 0.741 |
|  | SVM<sup>a</sup> | 0.968 | 0.969 | 0.967 | 0.813 |
|  | KNN | 0.883 | 0.908 | 0.884 | 0.295 |
|  | Naïve Bayes | 0.944 | 0.921 | 0.928 | 0.645 |
|  | Random forest | 0.803 | 0.895 | 0.846 | 0.000 |
|  | Logistic regression | 0.869 | 0.907 | 0.873 | 0.160 |

<sup>a</sup>Indicates the top three performing models.

The average precision-recall curve over all folds for each of the seven models for multi-class classification of the gram-negative and gram-positive bacteria in the LOPO cross-validation was plotted (**Fig. 4**). XGBoost had the largest AUC and was the outermost curve for both gram-negative and gram-positive bacteria, indicating the best average performance for multi-class classification of the expanded dataset. Individual precision-recall curves for each model are provided in the Supplementary Material for both gram-negative and gram-positive bacteria (**Supplementary Figs. S4 and S5**, respectively).

**Fig. 4.**
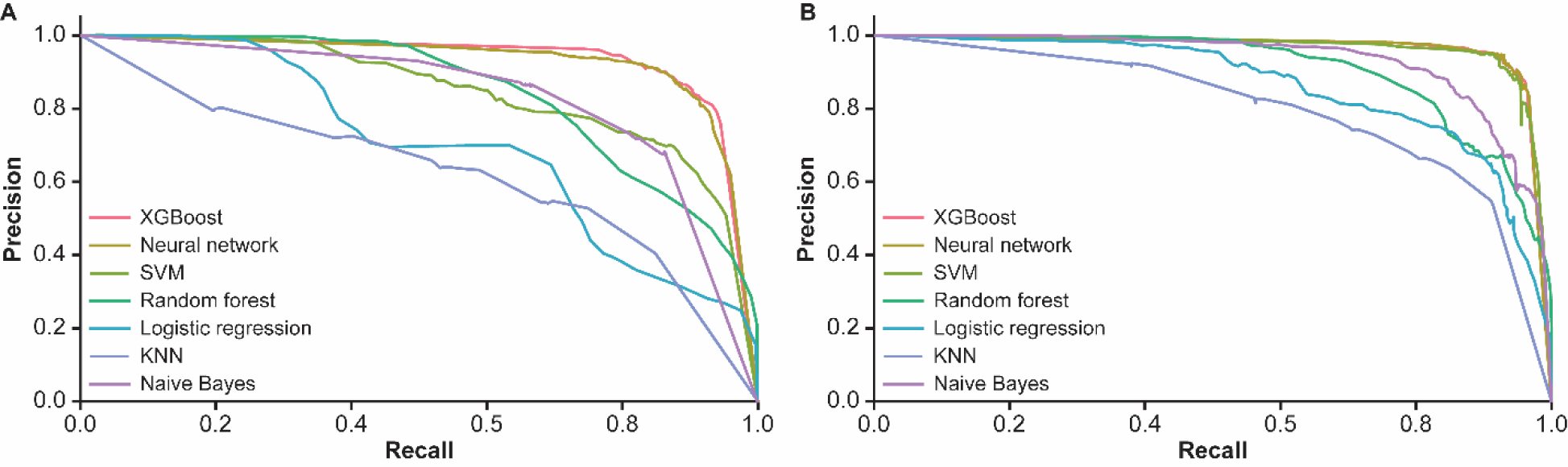
Average precision-recall curves for each of the ML models in LOPO cross-validation (multi-class classification) for (A) the 13 gram-negative bacteria and (B) 10 gram-positive bacteria.

Results from both cross-validation methods showed that neural network and XGBoost performed well with our pipeline in the case of gram-positive bacteria, with similar curves and a comparable AUC of 0.96. However, results from both cross-validation methods demonstrated XGBoost performed the best with our pipeline for gram-negative bacteria. Thus, this ML model was selected as the ideal option for our pipeline for further analyses on both gram-negative and gram-positive bacteria.

Accordingly, the nested 5FCV was run once again with our pipeline to finalize the hyperparameters, and the XGBoost model was trained on the expanded dataset. PSORTb was used as a benchmarking method to ensure that our pipeline made improvements in comparison to existing methods.

### 3.3. Benchmarking of mtx-COBRA

#### 3.3.1. Nested 5FCV

The expanded dataset was passed as input to PSORTb to facilitate comparison of results from the nested 5FCV conducted on the pipeline to PSORTb. On this dataset, predictions from PSORTb were compared with the ground truth, allowing calculation of the precision, recall, F1, and MCC scores, where the first three were adjusted for multi-class classification. These scores were used as the descriptors of the final performance of PSORTb on each dataset. Results obtained from the multi-class case were converted to the binary format and the scores were recalculated. In this last case, PSORTb was not retrained, as the original PSORTb code was not accessible; final multi-class results were converted to the binary case. However, because PSORTb was already trained and optimized, the results obtained from the conducted analyses were assumed comparable to the nested 5FCV completed on the pipeline. Thus, the final results on the complete database were comparable to the average results over the complete database of the nested 5FCV. For the gram-negative bacteria, the pipeline with the XGBoost ML model had higher precision, F1, and MCC scores than PSORTb, whereas PSORTb had a slightly higher recall score than the pipeline (**Table 5**). For both gram-negative and gram-positive bacteria in all other conditions, the pipeline with the XGBoost ML model performed better than PSORTb across all four classification metrics (**Tables 1 and 2**, respectively).

**Table 5.** Precision, recall, F1, and MCC scores for PSORTb tested on the expanded dataset and corresponding sores for mtx-COBRA with the XGBoost ML model.

| Bacteria | Classification | Cross-validation | Model | Precision | Recall | F1 score | MCC |
| --- | --- | --- | --- | --- | --- | --- | --- |
| Gram-negative | Multi-class | 5FCV | PSORTb | 0.947 | 0.979 | 0.962 | 0.733 |
|  |  |  | mtx-COBRA | 0.968 | 0.967 | 0.967 | 0.827 |
|  |  | LOPO | PSORTb | 0.920 | 0.801 | 0.849 | 0.734 |
|  |  |  | mtx-COBRA | 0.966 | 0.965 | 0.965 | 0.818 |
|  | Binary | 5FCV | PSORTb | 0.933 | 0.930 | 0.924 | 0.762 |
|  |  |  | mtx-COBRA | 0.950 | 0.951 | 0.950 | 0.837 |
|  |  | LOPO | PSORTb | 0.961 | 0.840 | 0.893 | 0.597 |
|  |  |  | mtx-COBRA | 0.945 | 0.941 | 0.939 | 0.797 |
| Gram-positive | Multi-class | 5FCV | PSORTb | 0.946 | 0.978 | 0.961 | 0.794 |
|  |  |  | mtx-COBRA | 0.981 | 0.981 | 0.981 | 0.895 |
|  |  | LOPO | PSORTb | 0.946 | 0.885 | 0.914 | 0.796 |
|  |  |  | mtx-COBRA | 0.968 | 0.966 | 0.967 | 0.857 |
|  | Binary | 5FV | PSORTb | 0.966 | 0.968 | 0.966 | 0.819 |
|  |  |  | mtx-COBRA | 0.969 | 0.970 | 0.969 | 0.832 |
|  |  | LOPO | PSORTb | 0.967 | 0.968 | 0.967 | 0.823 |
|  |  |  | mtx-COBRA | 0.967 | 0.967 | 0.965 | 0.800 |

#### 3.3.2. LOPO

PSORTb was run on the data for each of the 13 bacteria taken from the expanded dataset to facilitate comparison of the LOPO cross-validation method on the pipeline to the performance of PSORTb. Precision, recall, F1, and MCC scores of PSORTb on the data for each of the 13 bacteria were averaged for both multi-class and binary classifications. PSORTb was not retrained, and results were obtained using a comparable method where PSORTb was run on the data for each organism and results averaged. For the multi-class classification, the pipeline with the XGBoost ML model performed better than PSORTb for both gram-negative and gram-positive bacteria (**Table 5**). For the binary classification, the pipeline had higher recall, F1, and MCC scores than PSORTb for gram-negative bacteria, whereas the precision score of PSORTb was higher for gram-negative bacteria. For gram-positive bacteria, although results between the models were comparable, PSORTb had slightly higher recall, F1, and MCC scores, whereas the mtx-COBRA pipeline had the same precision as PSORTb.

#### 3.3.3. Test on held-out dataset

A final comparison between PSORTb and the pipeline was performed using the held-out dataset. This dataset comprised UniProt data for gram-negative bacteria, with all the homologs and sequences with greater than 30% similarity to those in the ePSORTdb and expanded datasets removed. Additionally, sequence clustering using MMseqs2 was conducted, creating a new dataset to be evaluated by PSORTb or the pipeline.

The held-out dataset was inputted to PSORTb, and the predictions obtained were compared with the ground truth to obtain precision, recall, F1, and MCC scores. Similarly, after being trained on the expanded dataset, the pipeline with the XGBoost ML model was run on the held-out dataset and the precision, recall, F1, and MCC scores were calculated. Because both XGBoost and neural network performed well for gram-negative and gram-positive bacteria, the pipeline with the neural network ML model was trained on the expanded dataset and tested on the held-out dataset to clarify which ML model performed the best with the pipeline.

The gram-negative bacteria precision, F1, and MCC scores for the pipeline with the XGBoost ML model were higher than those for PSORTb (**Table 6**). The recall score for the pipeline with the XGBoost ML model was comparable with the recall score for PSORTb, while a lower recall score was obtained for the pipeline with the neural network ML model (**Table 6**). PSORTb identified 195 proteins as having an “Unknown” SCL; the pipeline with the XGBoost ML model correctly predicted the SCLs for 164 (84.1%) of these proteins. The gram-positive bacteria precision, recall, F1, and MCC scores for the pipeline with the XGBoost ML model were higher than those for PSORTb as well as those for the pipeline with the neural network ML model (**Table 6**). PSORTb identified 223 proteins as having an “Unknown” SCL; the pipeline with the XGBoost ML model correctly predicted the SCLs for 166 (74.4%) of these proteins.

**Table 6.** Precision, recall, F1, and MCC scores for PSORTb and the pipeline with XGBoost and neural network ML models tested on the held-out dataset.

| Bacteria | Model | Precision | Recall | F1 score | MCC | Unknown |
| --- | --- | --- | --- | --- | --- | --- |
| **Gram-negative** | PSORTb | 0.925 | 0.981 | 0.950 | 0.647 | 195 unknown locations |
|  | XGBoost | 0.974 | 0.981 | 0.977 | 0.869 | 164/195 accurately predicted |
|  | Neural network | 0.969 | 0.975 | 0.972 | 0.846 | 158/195 accurately predicted |
| **Gram-positive** | PSORTb | 0.906 | 0.935 | 0.916 | 0.529 | 223 unknown locations |
|  | XGBoost | 0.951 | 0.965 | 0.957 | 0.759 | 166/223 accurately predicted |
|  | Neural network | 0.945 | 0.963 | 0.953 | 0.741 | 160/223 accurately predicted |

## 4. Discussion

Given that existing methods to determine protein SCLs are either experimental methods, which can be tedious and expensive, or bioinformatic methods, such as PSORTb, which returns many proteins with “Unknown” SCLs, our newly proposed bioinformatic pipeline, mtx-COBRA, aims to more accurately predict the SCL of proteins. We propose the use of a protein language model in combination with the XGBoost ML model to predict the SCL of proteins from gram-negative bacteria. The XGBoost model was chosen after analyses comparing multiple ML models with benchmarking factors.

Our pipeline is capable of multi-class classification and can identify the SCL of a protein in both gram-negative and gram-positive bacteria. By converting our multi-class classifications into a binary classification strategy, that is, by categorizing proteins as surface exposed or non– surface exposed, and then retraining and testing our model, we can judge the performance of our model as a binary classifier as well. After conducting analyses to determine the best training dataset, our pipeline was trained on an expanded dataset comprising data from the ePSORTdb and UniProt databases.

To benchmark the performance of our model and compare it to PSORTb, we used nested 5FCV and LOPO cross-validation methods. Our pipeline showed an improvement in F1 and MCC scores on the expanded and held-out datasets compared with PSORTb, regardless of bacteria type. The results from the gram-negative bacteria held-out dataset suggest that our model has significantly improved precision, F1, and MCC scores. In addition, our model accurately recovered the SCL for 84.1% of the proteins identified as “Unknown” by PSORTb. For the gram-positive bacteria, mtx-COBRA had improved precision, recall, F1, and MCC scores and recovered 74.4% of proteins identified as “Unknown” by PSORTb. Overall, our model improves the performance of SCL prediction of bacterial proteins in gram-negative and gram-positive bacteria compared with PSORTb, thus increasing the number of potential targets identified for antigen design in the vaccine development process.

## 5. Limitations

When considering the data spread for the LOPO cross-validation, all the ML models consistently underperformed for one gram-negative organism, *Sphingomonas paucimobilis*, and two gram-positive organisms, *Mycobacterium tuberculosis* and *Bacillus subtilisin.* It is noteworthy to point out the evolutionary distance on the respective taxonomic trees between *S. paucimobilis* and the other gram-negative bacteria investigated and that of *M. tuberculosis* and the other gram-positive bacteria included in this study (**Supplementary Fig. S1**). The slightly decreased performance on these evolutionarily distinct bacteria could be due to a lack of training data that contain organisms similar to or in a comparable location along the taxonomy tree as *S. paucimobilis* or *M. tuberculosis.* For *B. subtilis*, there could possibly be a lower number of proteins (in some or all SCLs) from bacteria that are similar to *B. subtilis* in the training dataset, causing a lower performance on this bacterium. However, the model included all organisms in the final training dataset, and this did not appear to hinder the overall performance of the model at successfully recovering the SCL for most of the proteins investigated, which were predicted as “Unknown” by PSORTb.

Another observation is that some ML models also underperformed for other organisms beyond those listed above. For example, the neural network underperformed for *Klebsiella pneumoniae* and *Citrobacter koseri,* despite their similarity to other organisms as depicted in the taxonomy tree. However, XGBoost managed to perform well on these organisms. This difference in performance could potentially be attributed to the difference in the ML models themselves, making XGBoost a better option for our pipeline than the other ML models.

Many companies use reverse vaccinology pipelines that involve an SCL prediction method to identify antigens that would be targeted during *in vitro* and *in vivo* immunogenicity testing [39]. An example of such a vaccine is the vaccine against *P. aeruginosa,* wherein reverse vaccinology was used to select antigens which were further tested *in vitro* and in mouse models [40]. Another vaccine, Bexsero, against meningococcal B, was developed using the reverse vaccinology process to identify the antigens to be included in the vaccine [41]. Bexsero has undergone clinical trials and is now commercially available to the public [42]. These examples show the importance of reverse vaccinology, which itself is not complete without SCL prediction.

Overall, the pipeline improves the accuracy of SCL prediction and increases the potential number of surface or extracellularly expressed antigens that can be considered for vaccine design. Given this improved performance, our method will assist with improving the accuracy of the reverse vaccinology process as a whole, expand the antigens that are targeted during *in vitro* and *in vivo* immunogenicity testing, and eventually assist with improving the quality and efficacy of the vaccines developed.

## 6. Conclusions

Our pipeline, mtx-COBRA, provides, on average, a more accurate method to predict SCL for proteins from gram-negative and gram-positive bacteria than the existing, widely used method of PSORTb. This is applicable to both the multi-class and binary classifications of SCL. The use of a protein language model provides an accessible and easy to use pipeline with greater efficiency to classify a larger number of proteins with currently “Unknown” SCLs than existing bioinformatic and experimental methods.

## Declaration of competing interest

At the time of this study, all authors were employed by Moderna, Inc., and may hold stock/stock options in the company.

## Acknowledgments

Medical writing and editorial assistance were provided by Ashlea İnan, PhD, Andy Kerr, PhD, and Audrey Shor, PhD, MPH, of MEDiSTRAVA in accordance with Good Publication Practice (GPP 2022) guidelines, funded by Moderna, Inc., and under the direction of the authors.

## Funding

This work was supported by Moderna, Inc. The funder was involved in the study design, collection, analysis, interpretation of data, and the writing of this article or the decision to submit it for publication.

## CRediT authorship contribution statement

Conceptualization: IA, MG, G-YC, EO; Formal analysis: IA; Funding Acquisition: N/A; Investigation: IA; Methodology: IA, AK, HA, EM, G-YC, EO; Project Administration: IA; Resources: N/A; Supervision: EO; Visualization: IA; Writing – original draft: IA; Writing – review & editing: IA, AK, HZ, MG, EM, G-YC, EO.

## Data sharing statement

Upon request, and subject to review, Moderna, Inc., will provide the data that support the findings of this study.

## Supplement

### Supplementary Tables

**Supplementary Table S1.** Gram-negative precision, recall, F1, and MCC scores for each model trained and tested on ePSORTdb using the 5FCV method.

| Classification | Model | Precision | Recall | F1 score | MCC |
| --- | --- | --- | --- | --- | --- |
| Multi-class | XGBoost<sup>a</sup> | 0.989 | 0.987 | 0.988 | 0.929 |
|  | Neural network<sup>a</sup> | 0.979 | 0.976 | 0.977 | 0.869 |
|  | SVM<sup>a</sup> | 0.977 | 0.965 | 0.971 | 0.831 |
|  | KNN | 0.972 | 0.964 | 0.968 | 0.820 |
|  | Naïve Bayes | 0.954 | 0.955 | 0.954 | 0.744 |
|  | Random forest | 0.957 | 0.912 | 0.925 | 0.650 |
|  | Logistic regression | 0.918 | 0.824 | 0.812 | 0.216 |
| Binary | XGBoost<sup>a</sup> | 0.979 | 0.979 | 0.979 | 0.918 |
|  | Neural network<sup>a</sup> | 0.964 | 0.965 | 0.964 | 0.863 |
|  | SVM<sup>a</sup> | 0.931 | 0.934 | 0.930 | 0.730 |
|  | KNN | 0.910 | 0.913 | 0.909 | 0.703 |
|  | Naïve Bayes | 0.887 | 0.886 | 0.886 | 0.635 |
|  | Random forest | 0.868 | 0.848 | 0.806 | 0.417 |
|  | Logistic regression | 0.846 | 0.810 | 0.727 | 0.095 |

<sup>a</sup>Indicates the top three performing models.

**Supplementary Table S2.** Gram-positive precision, recall, F1, and MCC scores for each model trained and tested on ePSORTdb using the 5FCV method.

| **Classification** | **Model** | **Precision** | **Recall** | **F1 score** | **MCC** |
| --- | --- | --- | --- | --- | --- |
| **Multi-class** | XGBoost<sup>a</sup> | 0.984 | 0.979 | 0.982 | 0.901 |
|  | Neural network<sup>a</sup> | 0.982 | 0.981 | 0.981 | 0.904 |
|  | SVM<sup>a</sup> | 0.981 | 0.979 | 0.980 | 0.894 |
|  | KNN | 0.972 | 0.951 | 0.960 | 0.829 |
|  | Naïve Bayes | 0.956 | 0.962 | 0.958 | 0.806 |
|  | Random forest | 0.974 | 0.930 | 0.946 | 0.790 |
|  | Logistic regression | 0.941 | 0.864 | 0.883 | 0.564 |
| **Binary** | XGBoost<sup>a</sup> | 0.979 | 0.979 | 0.979 | 0.917 |
|  | Neural network<sup>a</sup> | 0.974 | 0.974 | 0.974 | 0.896 |
|  | SVM<sup>a</sup> | 0.976 | 0.976 | 0.975 | 0.902 |
|  | KNN | 0.962 | 0.963 | 0.962 | 0.846 |
|  | Naïve Bayes | 0.961 | 0.958 | 0.959 | 0.842 |
|  | Random forest | 0.948 | 0.946 | 0.941 | 0.772 |
|  | Logistic regression | 0.883 | 0.888 | 0.864 | 0.458 |

<sup>a</sup>Indicates the top three performing models.

**Supplementary Table S3.** Gram-negative precision, recall, F1, and MCC scores for each model trained on ePSORTdb and tested on the expanded dataset using 5FCV method (multi-class classification).

| **Model** | **Precision** | **Recall** | **F1 score** | **MCC** |
| --- | --- | --- | --- | --- |
| XGBoost<sup>a</sup> | 0.976 | 0.979 | 0.977 | 0.883 |
| Neural network<sup>a</sup> | 0.962 | 0.965 | 0.963 | 0.807 |
| SVM<sup>a</sup> | 0.951 | 0.953 | 0.951 | 0.742 |
| KNN | 0.836 | 0.834 | 0.826 | 0.176 |
| Naïve Bayes | 0.922 | 0.931 | 0.925 | 0.622 |
| Random forest | 0.925 | 0.916 | 0.913 | 0.587 |
| Logistic regression | 0.853 | 0.831 | 0.809 | 0.189 |

<sup>a</sup>Indicates the top three performing models.

**Supplementary Table S4.** Gram-positive precision, recall, F1, and MCC scores for each model trained on ePSORTdb and tested on the expanded dataset using 5FCV method (multi-class classification).

| **Model** | **Precision** | **Recall** | **F1 score** | **MCC** |
| --- | --- | --- | --- | --- |
| XGBoost<sup>a</sup> | 0.967 | 0.977 | 0.971 | 0.882 |
| Neural network<sup>a</sup> | 0.964 | 0.972 | 0.968 | 0.864 |
| SVM<sup>a</sup> | 0.965 | 0.963 | 0.964 | 0.855 |
| KNN | 0.791 | 0.799 | 0.765 | 0.183 |
| Naïve Bayes | 0.926 | 0.926 | 0.930 | 0.707 |
| Random forest | 0.905 | 0.927 | 0.904 | 0.658 |
| Logistic regression | 0.827 | 0.818 | 0.771 | 0.309 |

<sup>a</sup>Indicates the top three performing models.

### Supplementary Figures

**Supplementary Fig. S1.**
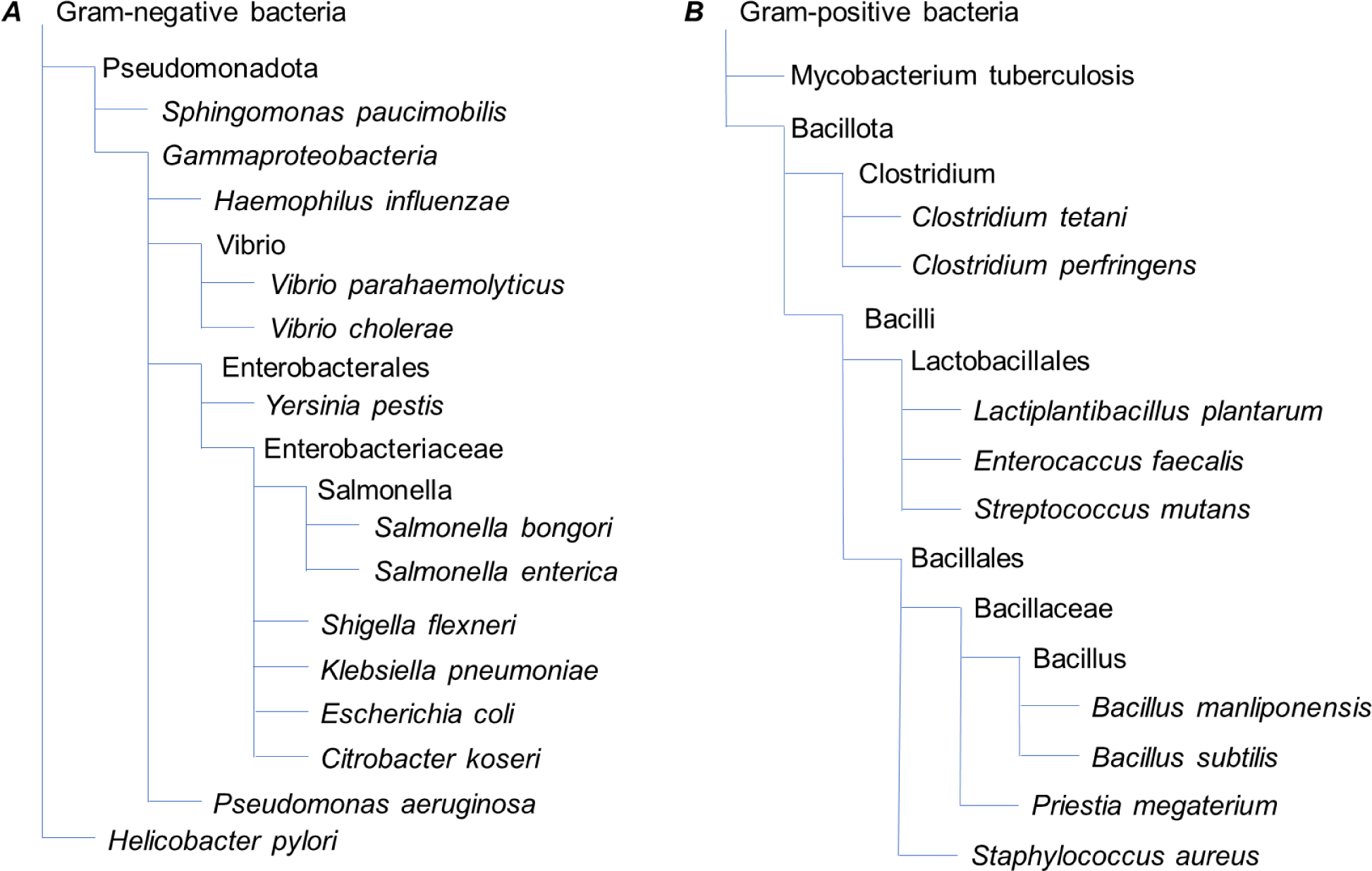
Taxonomic trees for the (A) gram-negative and (B) gram-positive bacteria chosen for LOPO cross-validation.

**Supplementary Fig. S2.**
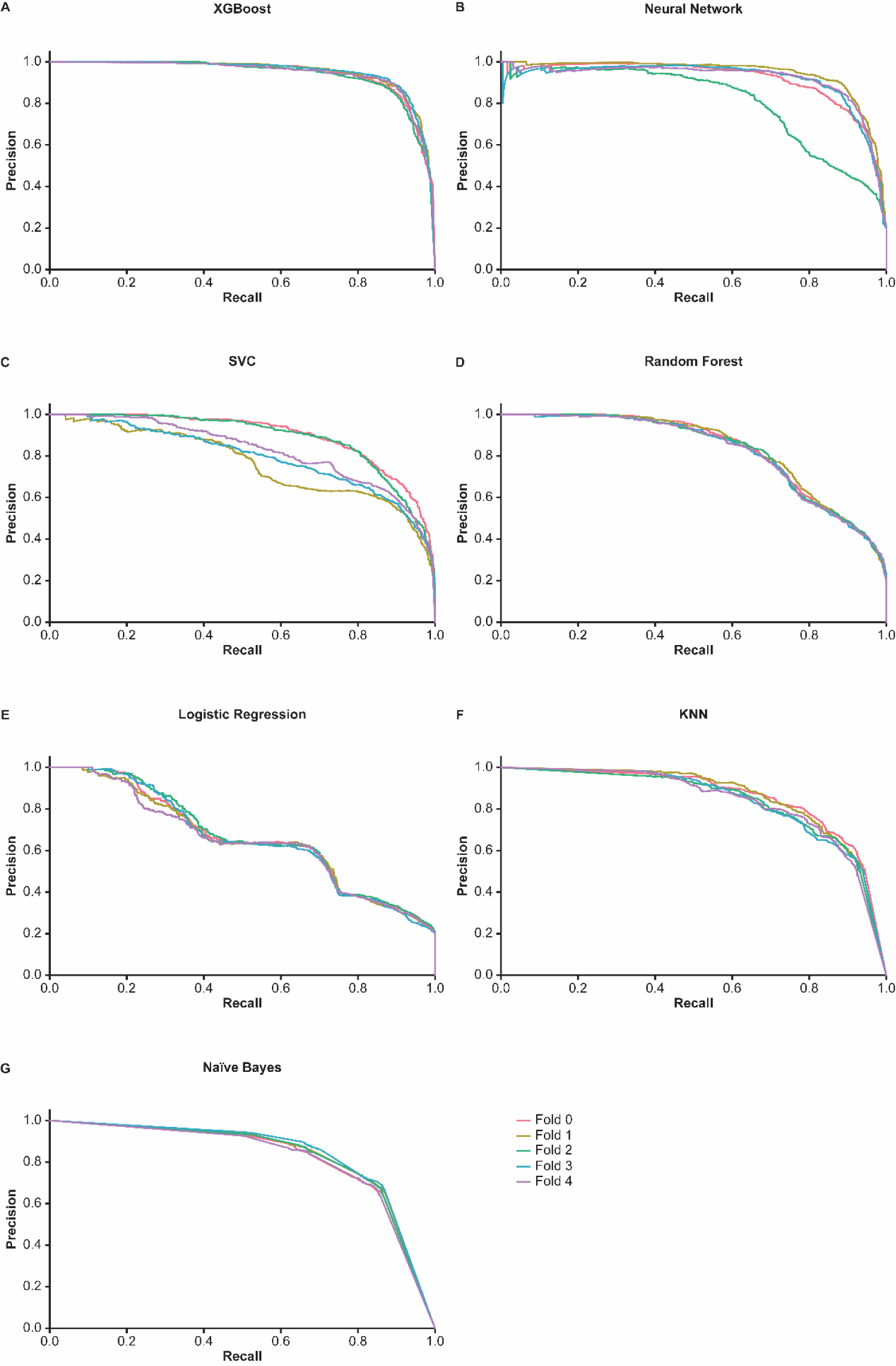
Gram-negative precision-recall curves for each fold and ML model in nested 5FCV (multi-class classification).

**Supplementary Fig. S3.**
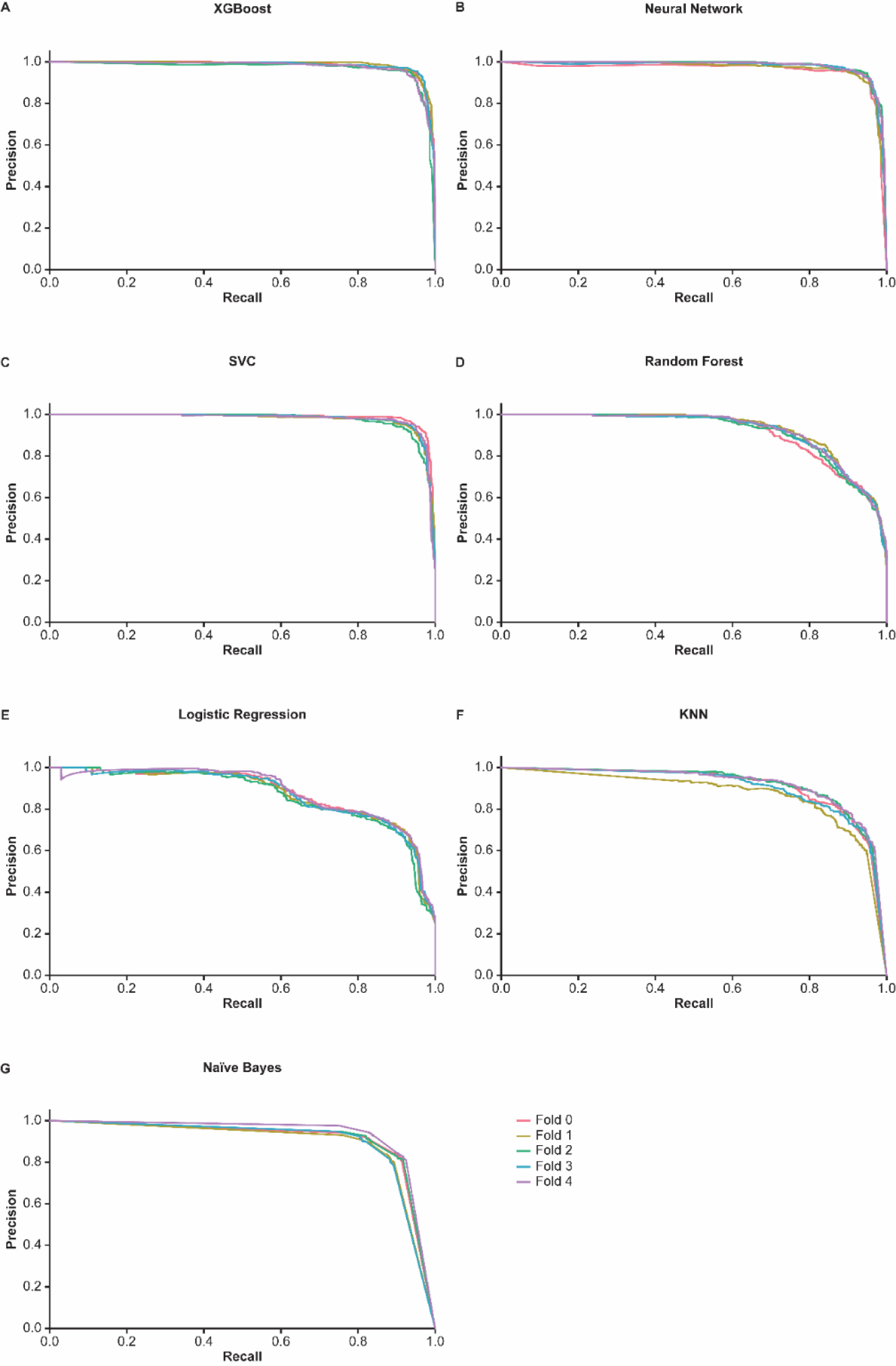
Gram-positive precision-recall curves for each fold and ML model in nested 5FCV (multi-class classification).

**Supplementary Fig. S4.**
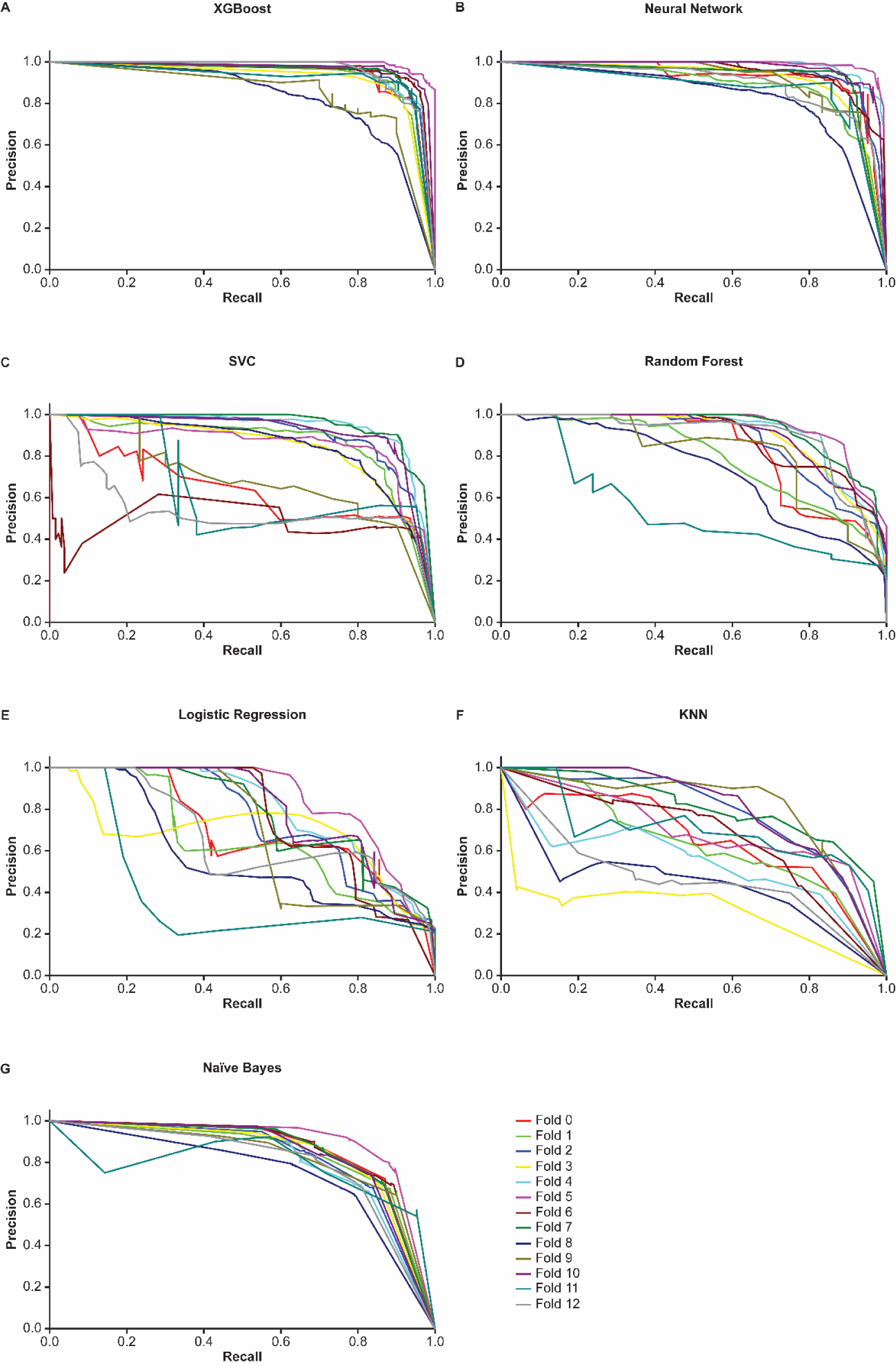
Gram-negative precision-recall curves for all ML models (multi-class classification).

**Supplementary Fig. S5.**
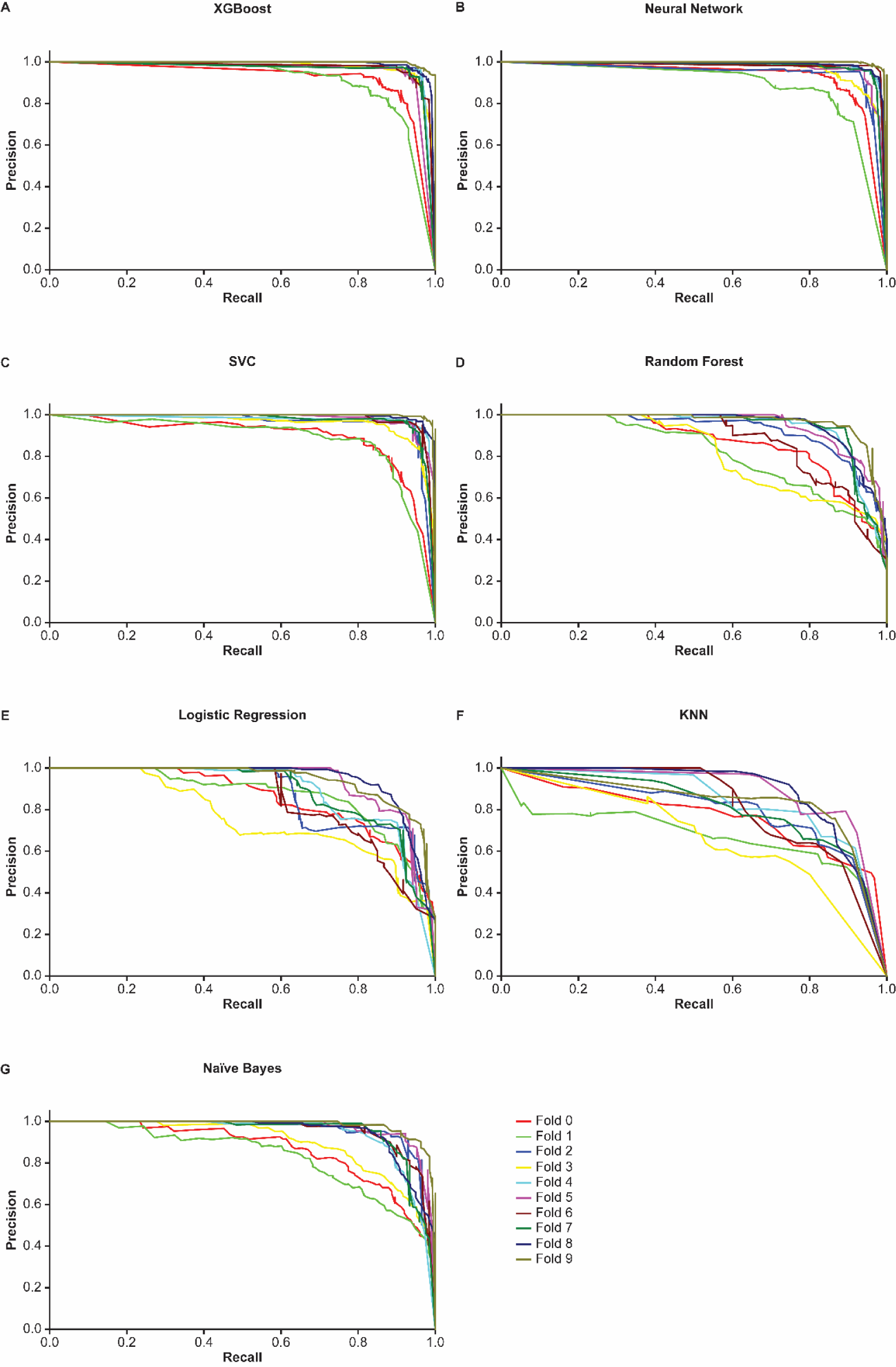
Gram-positive precision-recall curves for all ML models (multi-class classification).

